# Efficient generation of endogenous fluorescent reporters by Nested CRISPR in *Caenorhabditis elegans*

**DOI:** 10.1101/423970

**Authors:** Jeremy Vicencio, Adrian Martínez-Fernández, Xènia Serrat, Julián Cerón

## Abstract

CRISPR-based genome editing methods in model organisms are evolving at an extraordinary speed. Whereas the generation of deletion or missense mutants is quite straightforward, the production of endogenous fluorescent reporters is still inefficient. The use of plasmids with selection markers is an effective methodology, but often requires laborious and complicated cloning steps. We have established a cloning-free ribonucleoprotein-driven Nested CRISPR method that robustly produces endogenous fluorescent reporters. This methodology is based on the division of the GFP and mCherry sequences in three fragments. In the first step we use ssDNA donors (≤200 bp) to insert 5’ and 3’ fragments in the place of interest. In the second step, we use these sequences as homology regions for Homology Directed Repair (HDR) with a dsDNA donor (PCR product, ≈700 bp) including the middle fragment, thus completing the fluorescent protein sequence. This method is advantageous because the first step with ssDNA donors is known to be very efficient, and the second step, uses universal reagents, including validated PCR products and crRNAs, to create fluorescent reporters reaching reliable editing efficiencies as high as 40%. We have also used Nested CRISPR in a non-essential gene to produce a deletion mutant in the first step and a transcriptional reporter in the second step.

In the search of modifications to optimize the method, we tested synthetic sgRNAs, but we did not observe a significant increase in the efficacy compared to independently adding tracrRNA and crRNA to the injection mix. Conveniently, we also found that both steps of Nested CRISPR could be performed in a single injection. Finally, we discuss the utility of Nested CRISPR for targeted insertion of long DNA fragments in other systems and prospects of this method in the future.

## Introduction

The advent of CRISPR/Cas9 has allowed genetic engineering to progress at an unprecedented level. Naturally employed by bacteria as a defense mechanism, the Cas9 nuclease has been engineered to introduce blunt double-strand breaks (DSB) in target DNA when guided by an RNA duplex comprised of a generic trans-activating crRNA (tracrRNA) and a sequence specific CRISPR RNA (crRNA) (Mojica *et al*. 2005; Jinek *et al*. 2012). This cut is only made at sites complementary to the 20-nucleotide guide sequence within the crRNA in the presence of a downstream protospacer adjacent motif (PAM) site comprised of the bases 5’-NGG-3’ (in the case of *S. Pyogenes* Cas9). Thus, the ease of use and specificity of the technique has made it an attractive tool for genome editing in cellular systems and model organisms.

In the nematode *Caenorhabditis elegans*, gene editing is achieved via injection of a mix containing crRNA, tracrRNA, and Cas9 into the gonads. These components can be expressed from plasmids or added as independent molecules (commercially available) that form ribonucleoprotein complexes (RNPs) (Cho *et al*. 2013; Frokjaer-Jensen 2013; Waaijers *et al*. 2013). In the absence of a repair template, the non-homologous end joining (NHEJ) pathway is initiated, leaving behind deletions or small indels that are useful for generating non-specific mutations (Chen *et al*. 2013). However, when a repair template, in the form of a single-stranded oligonucleotide (ssODN) or double-stranded DNA with homology arms, is added into the mix, the homology-directed repair (HDR) pathway is initiated, allowing precise changes such as point mutations and defined deletions or insertions to be introduced into the genome (Paix *et al*. 2014). Simultaneous editing of the *dpy*-*10* locus leads to dumpy or roller phenotypes that facilitates rapid visual screening of genome-editing events, being commonly used as a positive control and co-marker (Kim *et al*. 2014; Arribere *et al*. 2014). A subset of *dpy*-*10* co-edited F_1_s would then be heterozygous for the edit of interest. One of the advantages of *C. elegans* is that it is hermaphroditic, and thus, self-fertilization will lead to ¼ of the F_2_ animals being homozygous for the desired edit. Coupled with the fast life cycle of this nematode (3-4 days at 20°C), producing the desired mutation takes only 10 to 15 days.

Despite the rate at which the optimization of the technique is progressing, it is not without limitations. These include variable efficiency of the sgRNA (Briner *et al*. 2014; Farboud and Meyer 2015; Liu *et al*. 2018), and potential off-target mutagenesis (Ran *et al*. 2013). In the case of insertions, another limiting factor is the length of commercially available ssODNs, which are commonly synthesized with a maximum length of 200 bp. In *C. elegans*, the homology arms need to be 35-45 bp long and therefore the maximum length of the new fragment that can be inserted by CRISPR with ssODNs is about 100-130 bp (Paix *et al*. 2014).

Insertions of this size are relatively easy to perform. However, the challenge currently lies in inserting larger fragments of DNA, such as the integration of fluorescent reporters into endogenous loci or gene replacements. Even though it has been demonstrated that this can be achieved by using targeting vectors with long homology arms (Dickinson *et al*. 2013; Schwartz and Jorgensen 2016; McDiarmid *et al*. 2018), plasmid construction usually involves cumbersome cloning steps. An alternative option is to use a PCR product with 35-bp flanking sites that are homologous to the insertion site as a repair template (Paix *et al*. 2015). However, in our hands, and in the experience of other colleagues, the efficiency of this method is very low (Dokshin *et al*. 2018).

In order to overcome this challenge, we developed a Nested CRISPR protocol that can consistently generate protein::EGFP or protein::mCherry fusions without the need for cloning. It involves the insertion of the fluorescent protein of interest in two steps. The first step involves a ≈120-bp in-frame insertion, at the N-terminus or C-terminus of the target gene, consisting of the joint 5’ and 3’ ends of EGFP or mCherry using an ssODN as a repair template. This fragment contains a new PAM site and protospacer sequence that, in a second step, allow the in-frame insertion of the remaining sequence of about 700 bp, depending on the number and length of introns included in the donor, using a universal PCR product as a repair template. We demonstrate that the high editing efficiency in the first step is not limiting (≈15-69%), and that the remaining larger fragment can be inserted in the second step with ≈7-40% of efficiency. As a result, we present data for EGFP and/or mCherry integration across five genes, including seven protein fusions and one transcriptional reporter, demonstrating the efficacy of the Nested CRISPR method.

## Results

### Endogenous translational fluorescent reporters by Nested CRISPR

We were interested in making an endogenous fluorescent reporter for *prpf*-*4.* At that time, the most straightforward method was the use of CRISPR/Cas9 ribonucleoprotein complexes and a PCR product with 35 bp homology arms as donor (Paix *et al*. 2015). We obtained a single positive event after four experiments consisting of 72 injections and 366 PCRs (0.3% of efficacy). We used the same methodology for five more genes, but we failed to generate other reporters after many injections. Thus, we decided to investigate an alternative cloning-free method to generate endogenous fluorescent reporters in *C. elegans*. Since the targeted insertion of long double-stranded DNA (dsDNA) templates was much more difficult than genome editing with small (≤200 nt) single-stranded DNA, we reasoned that a nested approach that always uses the same, previously validated, dsDNA and crRNA to insert the longer fragment would be an efficient method to produce endogenous EGFP and mCherry reporters. Thus, we designed a pipeline to produce homozygous translational reporters in five generations (approximately three weeks in *C. elegans*) (**Figure 1**). This pipeline works as follows: in the case of the Enhanced Green Fluorescent Protein (EGFP), we divide the 866 nt of the EGFP sequence, including three introns, in three sequences of 58 nt + 752 nt + 56 nt that are designated as fragments 1, 2 and 3 respectively (**Figure 2A**). Using a crRNA specific for the targeted gene, we can insert the block of sequences 1 and 3 in-frame in the place of interest by using single stranded oligodeoxynucleotides (ssODNs) as donors (**Table S1, S2**). As positive control, we use the co-CRISPR strategy (Kim *et al*. 2014; Arribere *et al*. 2014). Thus, only plates with the presence of *dpy*-*10*-edited worms, either dumpy or roller animals, are screened for insertions by PCR (**Figure 1**). Once we have homozygous animals for the 1-3 block, we sequence the insertion to ensure that the 1-3 fragment is in frame. We detected a right sequence in 62% of the cases (13 out of 21) and therefore we recommend to sequence three positives (**Table 1**).

**Figure 1.**
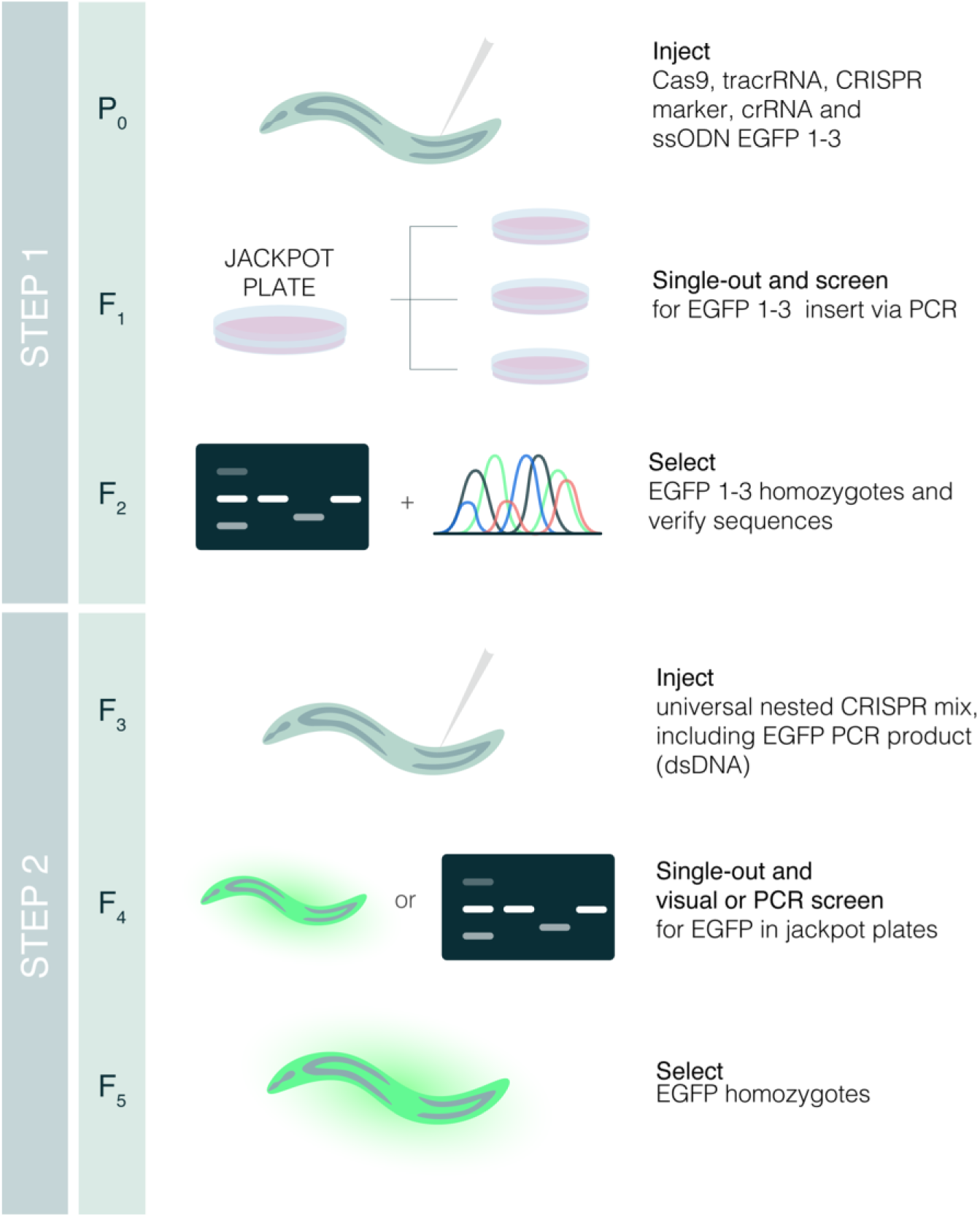
Nested CRISPR pipeline. Young adult animals are injected in the germline with a mix containing Cas9, tracrRNA, crRNA and ssODN repair template for the fluorescent protein of interest (EGFP in this example), and a marker of CRISPR efficiency (crRNA and ssODN for *dpy*-*10(cn64)* in our case as in Arribere et al, 2014). Injected animals are singled-out in NGM plates. After 3-4 days, animals from plates with positives for the CRIPSR marker (jackpot plates) are singled out in NGM plates and genotyped for the insertion of interest once they laid progeny. The progeny of F_1_ animals positive for the insertion is singled out to obtain F_2_ homozygous animals, and insertions verified by Sanger sequencing. Bands in gel and sequencing peaks are illustrative. In the second step, F_3_ young adults with the insertion of fragment 1 and 3 in frame are injected with a universal mix containing Cas9, tracrRNA, crRNA for fragment 1-3, dsDNA (PCR product of the fluorophore of interest) as repair template, and CRISPR marker. The fluorescence could be observed in F_4_ positive animals. Otherwise, a PCR screen for positives among F_4_ animals from jackpot plates is required. F_4_ animals heterozygous for the insertion will give rise to F_5_ homozygotes for the translational reporter of interest.

**Figure 2.**
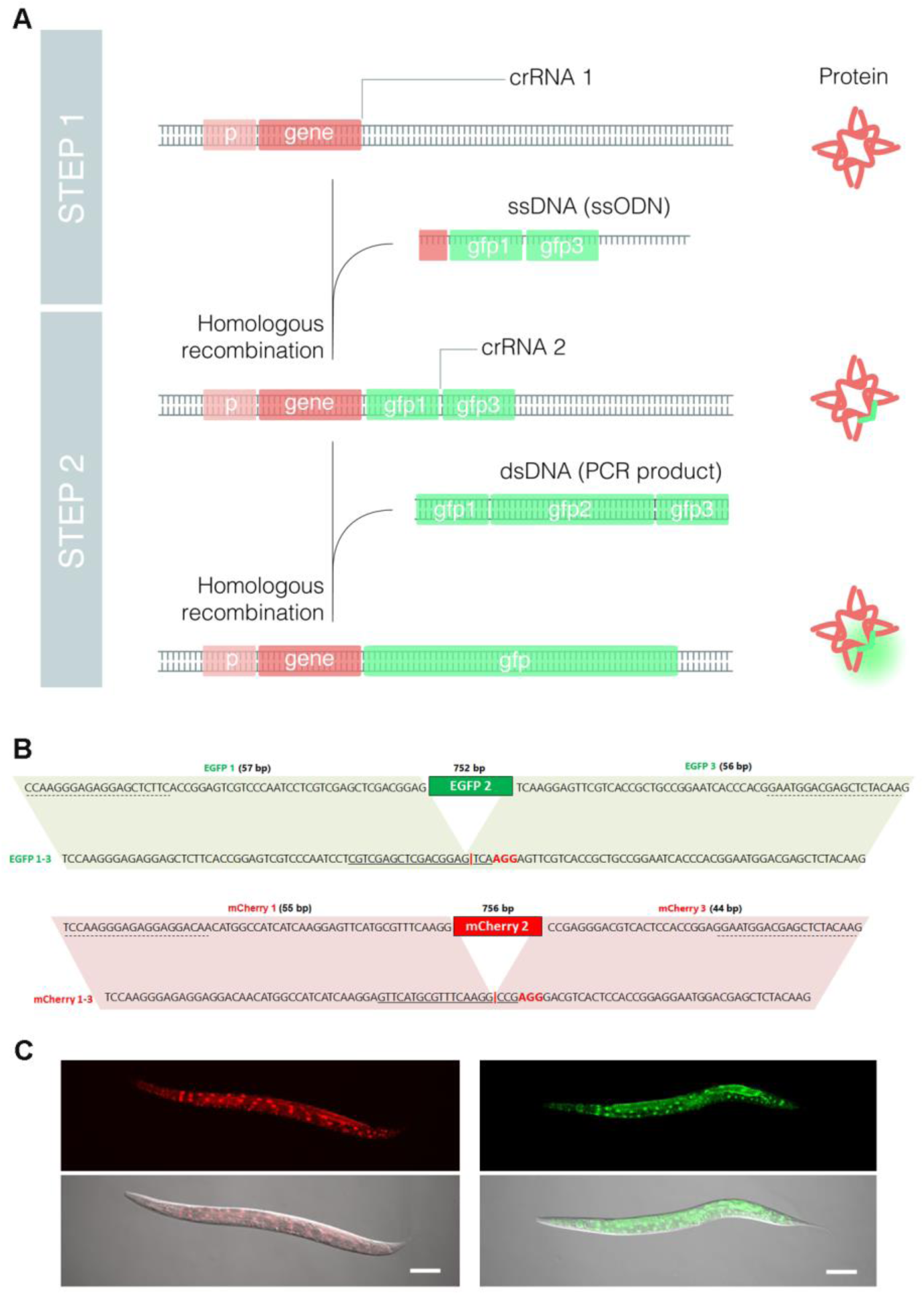
Translational endogenous reporters by Nested CRISPR. **(A)** Scheme of molecular events to generate a translational fluorescent reporter (GFP in this case) by Nested CRISPR. A gene-specific crRNA (crRNA 1) is required to assemble Cas9 RNP complexes that cut at the 5’ or 3’ end of the gene. Along with these RNPs, the injection mix contains an ssODN with two homology arms of 35-45 bp (depending on the distance from the cut site) that is inserted in the place of interest by homologous recombination. In the second step, RNPs contain a universal crRNA (crRNA 2) that cut the gfp 1-3 specific targeted sequence or protospacer. Then a universal dsDNA molecule, resulting from PCR amplification of GFP, is used as repair template to generate a translational reporter. **(B)** Details of sequences and homology regions of EGFP and mCherry for the universal Step 2. The nucleotides at the bottom represent the sequences of fragments 1-3 for EGFP and mCherry after Step 1. Solid lines correspond to the 20-nt guide sequence and nucleotides in red represent the PAM sequence. Targeted sequences in EGFP and mCherry result from the fusion of native fragments 1 and 3, without the need to change any nucleotide. The red vertical bar represents the cut site. At the top is the sequence of the dsDNA repair template with homology with fragments 1-3. Primer annealing sites for PCR amplification of EGFP and mCherry are labeled with dashed lines. Parallelograms mark homology regions. **(C)** Representative pictures of *prpf*-*4*::mCherry and *prpf*-*4*::EGFP translational reporters generated by Nested CRISPR. Scale bars equivalent to 100 μm.

**Table 1.** Summary of Nested CRISPR experiments.

| Experiment | Plates with<br>*dpy-10* edits/<br>injected worms | positives<sup>A</sup>/worms<br>screened (%) | In-frame<br>sequences<sup>D</sup> | Cas9<br>concentration<br>(ng/μl) |
| --- | --- | --- | --- | --- |
| *prpf-4*::EGFP Step 1 | 1/13 | 6/23 (26.09) | 2 of 2 | 1500 |
| *prpf-4*::EGFP Step 2 | 7/23 | 13/32 (40.63) | 13 of 13 | 1640 |
| *prpf-4*::EGFP Step 2 syn-sgRNA | 4/20 | 2/24 (8.33)<sup>E</sup> | N/A | 250 |
| *prpf-4*::mCherry Step 1 | 6/12 | 3/23 (13.04) | 2 of 3 | 250 |
| *prpf-4*::mCherry Step 2 | 11/28 | 16/43 (37.21) | 16 of 16 | 250 |
| *prpf-4*::mCherry Step 2 syn-sgRNA | 7/28 | 26/108 (24.07) | 3 of 3 | 250 |
| *prpf-4*::mCherry single injection | 11/29 | 16/77 (20.78)<sup>E</sup> | N/A | 250 |
| *gtbp-1*::EGFP Step 1 | 9/14 | 3/24 (12.50)<sup>B</sup> | 1 of 3 | 1000 |
| *gtbp-1*::EGFP Step 2 | 9/24 | 10/46 (21.74) | 10 of 10 | 250 |
| *gtbp-1*::EGFP Step 2 syn-sgRNA | 4/13 | 3/34 (8.82) | 3 of 3 | 250 |
| *gtbp-1*::mCherry Step 1 | 6/15 | 22/32 (68.75) | 1 of 3 | 250 |
| *gtbp-1*::mCherry Step 2 | 7/18 | 7/62 (11.29) | 7 of 7 | 250 |
| *pgl-1*::EGFP Step 1 | 5/24 | 7/25 (28.00)<sup>C</sup> | 1 of 1 | 1000 |
| *pgl-1*::EGFP Step 2 | 5/16 | 1/13 (7.69) | 1 of 1 | 250 |
| *pgl-1*::mCherry Step 1 | 3/13 | 24/45 (53.33) | 2 of 5 | 250 |
| *pgl-1*::mCherry Step 2 | 4/18 | 14/39 (35.89) | 7 of 14 | 250 |
| K12C11.3p::mCherry Step 1 | 7/37 | 21/56 (37.50) | 3 of 3 | 250 |
| K12C11.3p::mCherry Step 2 | 4/14 | 3/35 (8.57) | 3 of 3 | 250 |
| *ubh-4*::EGFP Step 1 | 2/8 | 8/15 (53.30)<sup>C</sup> | 1 of 1 | 500 |
| *ubh-4*::EGFP Step 2 | 9/27 | 4/39 (10.26)<sup>E</sup> | N/A | 250 |

<sup>A</sup> Based on PCR genotyping (amplicons of the correct size are considered positives regardless of whether or not the insertion is in-frame)
<sup>B</sup> Pools of 2-3 worms
<sup>C</sup> Pools of 2 worms
<sup>D</sup> In-frame sequences represent the fraction of worms that are confirmed by Sanger sequencing (Step 1) or by visual screening (Step 2) over the number of positives by PCR genotyping.
<sup>E</sup> Identification of positives via visual screening only.

Then, we use sequences of EGFP 1 and 3 as homology regions to insert the remaining EGFP sequence by using a EGFP 1-3 specific crRNA and a PCR product amplified from the pJJR82 EGFP plasmid (**Figure 2B, Table S3, Video S1**). Importantly, the injection mix in this second step is universal, using the same crRNA and PCR product, increasing reproducibility and reducing costs (**Table S3**). Scoring of positives in the progeny can be visual if the fluorescence signal and the stereoscope, or microscope, are adequate. Otherwise, it is necessary to perform a PCR-based screen among animals, individuals or pools, from *dpy*-*10* edited plates.

We first applied the Nested CRISPR pipeline to *prpf*-*4* to produce an EGFP reporter. In the first step, we injected 13 animals and genotyped 24 F_1_ worms obtaining 26% of positives. In the second step, we injected 23 animals and genotyped 32 F_1_ worms obtaining 40% of positives. Whereas in the past we spent months to generate one strain, following the Nested CRISPR pipeline we obtained 13 lines for *prpf*-*4*::EGFP in three weeks (**Figure 2C**). Thus, we decided to validate and consolidate this methodology. Then we generated a *prpf*-*4*::mCherry reporter with similar efficiency (**Figure 2B, 2C**) (**Table 1**). Next, we made endogenous EGFP and mCherry reporters for other genes, *gtbp*-*1* and *pgl*-*1*, that were previously used to test CRISPR/Cas9 methods (Paix et al., 2015; 2016). Efficacies ranged from 12% to 70% in the first step, and from 7% to 37% in the second step (**Figure 3**) (**Table 1**). Finally, we attempted to create a translational reporter for *ubh*-*4*, a gene that had not previously been edited by CRISPR. We obtained a 53% of efficacy in the first step, and 10% in the second step.

**Figure 3.**
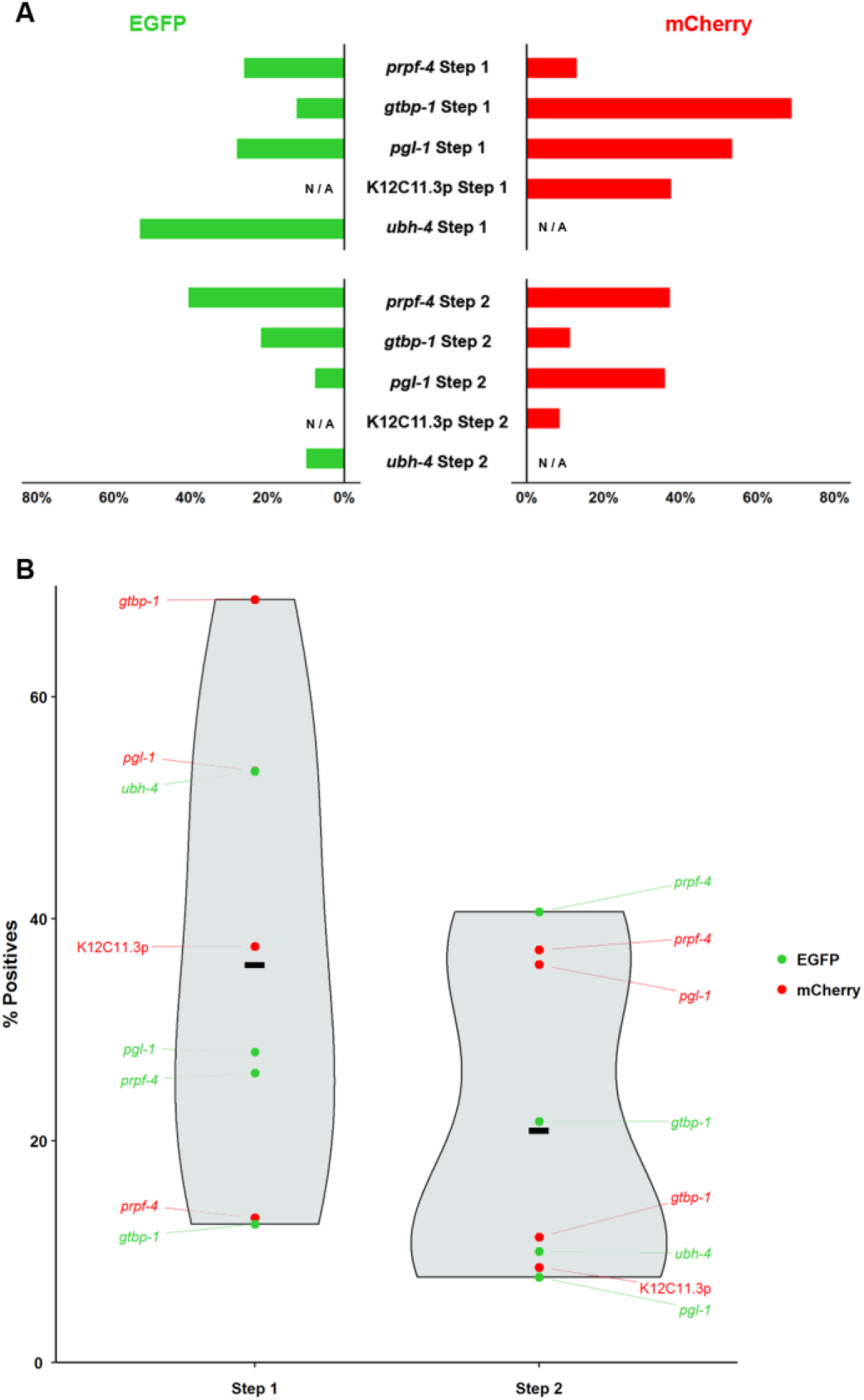
Efficiency of Nested CRISPR experiments. **(A)** Horizontal bar plot representing the efficiency of each experiment, as described in Table 1. N/A = not applicable. **(B)** Graph showing the distribution of efficacy values represented in the previous panel. Violin plot (in blue) illustrates the kernel probability density of each step’s efficiencies. Red and green dots represent efficiencies (% of positives) of each attempt to introduce mCherry or EGFP, respectively. Black lines represent the mean efficiency in each step.

As stated above, a Nested CRISPR universal mix can be used for all Step 2 injections to increase the reproducibility of Homology Directed Repair (HDR) efficiency. Still, we observed variability in the efficiency of step 2 (**Figure 3B**) that could be due to local chromatin structure or other factors that we do not yet understand. Still, the relevant point is that, differently from the past, we succeeded to produce several lines in all our attempts to generate endogenous fluorescent reporters.

### Nested CRISPR pipeline to generate a deletion mutant and a transcriptional reporter

In our experience, removing the whole ORF of a given gene by CRISPR using two crRNAs at the 5’ and 3’ ends (Chen *et al*. 2014) works very well with the addition to the injection mix of an ssODN as repair template, producing mutants with a precise deletion.

We reasoned that in the case of non-essential genes, which are about 80-85% of the *C. elegans* genome (Kemphues 2005), our Nested CRISPR method could be used to produce both a deletion mutant and a transcriptional reporter in the same pipeline (**Figure 4, Video S2**). We tested this approach in the gene K12C11.3, which encodes a non-essential copper transporter. First, we made a 1339 bp deletion that removes most of the K12C11.3 coding sequence. We used two crRNAs to cut just after the initiation codon and right before the stop codon, and an ssODN donor containing the mCherry 1-3 fragment with two 35 bp homology arms (with homology upstream of the initiation codon, and downstream of the stop codon). In this first step we obtained 37% of positive events after the PCR screen. In the second step, we injected 35 animals and produced three transcriptional reporters for K12C11.3 (8% of efficacy) (**Table 1**). Therefore, nested CRISPR is an advantageous option when generating deletion mutants of non-essential genes as the knockout obtained is already primed for the generation of a transcriptional reporter.

**Figure 4.**
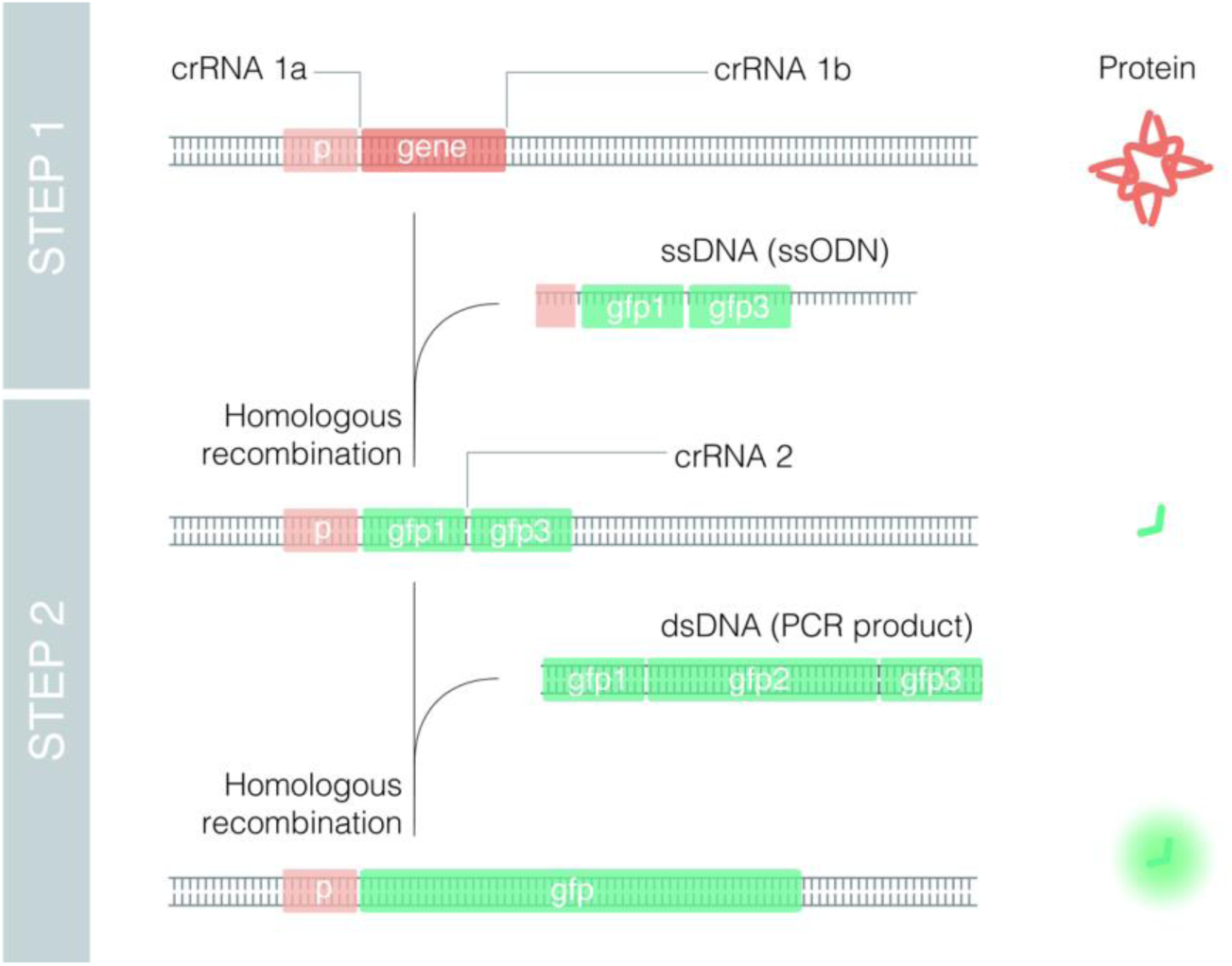
Deletions and transcriptional endogenous reporters by Nested CRISPR. Scheme of molecular events to generate a deletion and a transcriptional fluorescent reporter (GFP in this case) by Nested CRISPR. Two gene-specific crRNA (crRNA 1a and crRNA 1b) and a ssODN donor are required to produce a deletion of the gene and an in-frame insertion of fragment 1-3. In the second step, RNPs contain a universal crRNA (crRNA 2) that cut the gfp 1-3 specific targeted sequence. Then a universal dsDNA molecule, resulting from PCR amplification of GFP, is used as repair template to generate a transcriptional reporter.

### Synthetic sgRNAs are functional

In the use of RNPs for CRISPR, researchers have been using crRNA:tracrRNA duplexes that are bought separately and subsequently mixed 1:1 later to make RNPs with Cas9. Recently, synthetic sgRNAs become commercially available (IDT). This new synthetic sgRNA consists of a single RNA oligonucleotide containing both the target-specific crRNA and the universal tracrRNA. Synthetic sgRNAs contain chemical modifications that may improve the stability and we wondered if they would increase the efficacy of CRISPR in *C. elegans.* We found that synthetic sgRNAs work correctly but we did not observe a clear improvement in the efficacy (**Table 1**). However, synthetic sgRNAs consumes less volume in the injection mix compared to adding crRNA and tracrRNA independently, and this could be convenient in cases and/or systems where final volume of the mix is critical.

### Two steps in one injection

Theoretically, the two steps of Nested CRISPR could occur in the same animal by co-injecting all crRNAs and donors (gene-specific for Step 1 and universal for Step 2). Since the universal step 2 crRNA and the gene-specific step 1 EGFP or mCherry 1-3 ssODN are non-complementary, Cas9 will only recognize the step 2 protospacer sequence after the strand complementary to the ssODN repair template has been synthesized via HDR. Thus, it is said that Cas9 will initiate the cut for step 2 only when step 1 has been completed. We injected 29 animals and obtained 11 plates with *dpy*-*10*-edited animals. Out of 77 *dpy*-*10*-edited animals we observed 16 positives for *prpf*-*4*::mCherry, which corresponds to an efficiency of 20% (**Table 1**).

## Discussion

CRISPR technology is evolving rapidly and such speed does not facilitate the consolidation of protocols through reproducibility in distinct laboratories. Thus, an efficacy of 16% to insert two loxP sites to produce conditional KOs in mice (Yang *et al*. 2013), has recently been proved to be actually close to 1% by the mice community (Gurumurthy *et al*. 2018). In *C. elegans*, the CRISPR-based insertion of fluorescent tags using PCR products with 35 bp homology arms as donors does not seem to be as efficient as originally reported (Paix *et al*. 2015). Such discrepancy has formally been mentioned for the first time in a recent publication (Dokshin *et al*. 2018). These inconsistencies between efficacies are probably due to factors that are still unknown because of the speed at which the field is progressing. Some labs use in-house purified Cas9 whereas other labs use commercial Cas9, and each lab has different equipment to perform microinjections in the worm germline. These differences are a source of variability. Unfortunately, differently from the mice community, there is no coordinated effort to determine the real efficiency of the different techniques in *C. elegans*.

CRISPR/Cas9 reagents are now commercially available and the cost is affordable for most labs. Thus, the use of common reagents should help to unifying the outcome of CRISPR methods. However, the way to perform microinjection in *C. elegans* varies from lab to lab, and from person to person, and this could also be source of variability in the amount of Cas9 injected in the germline. Since high concentration of Cas9 appears to be toxic (Dokshin *et al*. 2018), microinjection could be considered a critical point. In fact, there is quite variability in the efficacy among worms injected in the same experiment by the same person.

### Present and future of Nested CRISPR

This Nested CRISPR approach has several advantages (**Table 2**) and has changed our experimental life in terms of generating endogenous reporters. While we previously spent months performing CRISPR experiments attempting to generate reporters of six distinct genes and being successful only once, following the Nested CRISPR method we have succeeded in all eleven attempts to make endogenous fluorescent reporters (**Table 1**). Other researchers should reproduce our success rate with Nested CRISPR since all the elements that we use are commercially available, and the reagents and conditions for the second step are universal. This methodology is feasible for researchers with limited knowledge of molecular cloning and will facilitate the production of endogenous reporters to study the real expression of a given gene. Therefore, expression patterns inferred from extrachromosomal reporters or molecular constructs inserted randomly in the genome, and often as multicopy, should be revised and validated.

**Table 2.** Advantages of the NESTED CRISPR method.

|  |
| --- |
| All ingredients are commercially available (except for universal EGFP or mCherry PCR products which can be easily prepared). |
| The homogeneity of the reagents allows inter-laboratory comparative analysis of CRISPR efficiency. |
| The second step utilizes the same reagents and conditions in all cases. Thus, it is reliable and the difficulty of inserting long dsDNA fragments is bypassed. |
| Given the universality of the second step, it is scalable to genome-wide projects. |
| Different EGFP/mCherry variants can be used, with distinct introns that may influence expression levels and silencing of exogenous DNA. |
| The fluorescent protein PCR product repair templates amplified with high-fidelity polymerase are universal, thus avoiding multiple PCRs with expensive polymerases. |
| An insertion in the first step indicates an accessible chromatin environment for a cut in the second step. |

We have used the co-CRISPR system as a marker of valid injections. However, we have not found a strict correlation between positives and plates with a high number of *dpy*-*10* edited animals (**Figure S1**). Alternatively, other co-injection markers as extrachromosomal plasmids harboring fluorescent reporters, antibiotic resistance genes, or dominant alleles can be used (Norris *et al*. 2015; Prior *et al*. 2017; Dokshin *et al*. 2018). However, the use of plasmids could be a source of variability (mutations, distinct DNA preparations, etc.) between labs that can be bypassed by using the same crRNA for *dpy*-*10.* Still, to date, it is impossible to predict which injected animals will produce more positives.

Since the structure of the guide RNA can influence the activity of Cas9 (Lim *et al*. 2016), we tested synthetic sgRNA for some injections and we did not observe an improvement compared to injecting tracrRNA and crRNA separately. A positive aspect of synthetic sgRNAs is that the volume of the injection mix can be reduced and a potential competition between two crRNAs for the tracrRNA can be avoided.

We also tried universal GFP and mCherry megamers (long ssDNA) plus two small ssODNs as homology bridges to induce *in vivo* assembly of linear DNAs (Paix *et al*. 2016), but we only obtained partial insertions. Although the price of megamers is still high and the synthesis of long ssDNA can present difficulties, the use of megamers in CRISPR should be explored and evaluated in the future (Quadros *et al*. 2017).

The race to develop more efficient and reliable CRISPR methodologies has not stopped. *C. elegans* researchers have found several factors that can influence the efficacy of CRISPR. As examples, the orientation of the ssDNA repair seems to influence the efficiency of insertion (Katic, Xu, & Ciosk, 2015), certain nucleotides before the PAM sequence could be more convenient for Cas9 cutting (Farboud and Meyer 2015), and new variants of Cas9 protein may be more specific and efficient (Bell *et al*. 2016). Despite these, the insertion of long DNA fragments has a limited efficiency. Recently, two studies have proposed modifications in dsDNA templates to improve efficacy of genome editing. These studies suggest the use of 5’ modifications in the dsDNA donors and the use of hybrid PCR products with 120 bp of ssDNA overhangs (Ghanta *et al*. 2018; Dokshin *et al*. 2018). These alternatives are attractive but can be costly, particularly for large-scale projects. Thus, the generation of hybrid PCR products requires long primers (140 nt) that may need to be optimized whereas the PCR for the second step in Nested CRISPR is already optimized and the product can be reused for several experiments. In any case, these studies and ours are open to the community and will certainly help different labs in finding the convenient methodology that suits their expertise and resources. In fact, a coordinated effort from the *C. elegans* community is necessary to compare distinct approaches for inserting long DNA fragments by CRISPR. Meanwhile, the onset of different but efficient CRISPR methodologies will facilitate the widespread generation of endogenous reporters that will step up many research projects.

### Nested CRISPR is scalable

We used Nested CRISPR to generate a deletion mutant and a transcriptional reporter in the same pipeline. This is a strategy that could be considered for large-scale projects because at the time that a collection of deletion mutants is made, these strains are ready for a universal second step that generate transcriptional reporters. Conveniently, the number, length, and sequences of introns can be modified in the second step. This is relevant because the number and length of introns can influence the transcriptional rate, and the sequence of these introns can influence germline silencing (Frokjaer-Jensen 2013; Heyn *et al*. 2015). Thus, we found the modular and flexible nature of the second step of Nested CRISPR of great value.

The fact that we used homozygous animals with the 1-3 fragment in our deletion plus transcriptional reporter pipeline is a handicap for essential genes (approximately 20% of genes) whose deletions need to be maintained as heterozygous strains. However, such difficulty can be bypassed by doing Nested CRISPR in a single injection, which is feasible as we observed in a pilot assay (Table 1).

Recently, a scalable strategy to create mutants in *C. elegans* has been suggested. Such strategy relies in the insertion of an ssODN with STOP codons in the three different reading frames (Wang *et al*. 2018). This is a smart approach but, beside the concern of having some residual translation due to an inefficient Non-sense Mediated Decay (NMD), using Nested CRISPR results in a deletion mutant strain ready to later produce a fluorescent reporter.

### Universality of Nested CRISPR

The efficacy of Nested CRISPR relies in the use of a universal and reliable step to insert long fragments of DNA. Perhaps, the insertion of a long piece of DNA in a genomic region that has previously been edited (step 1) could be facilitated. It is known that chromatin state influences CRISPR-Cas9 editing efficiencies (Verkuijl and Rots 2018) and it is possible that the first cut makes the chromatin more accessible or sensitive for an subsequent cut. It any case, the cut required for the first step of Nested CRIPSR indicates that the chromatin in that region is accessible for Cas9 to perform the second and more limiting step.

The mechanisms of CRISPR/Cas9 genome-editing in distinct organisms seem to be very similar and therefore, any technical advance in *C. elegans* could be applied to other models. In the case of Nested CRISPR, the need for five generations to obtain homozygous animals could be a handicap in other animals with longer life cycles. Still, if somebody plans to make a deletion mutant for a given gene, it will make sense to make it ready for a second step that allows the generation of an endogenous fluorescent reporter. We have performed one pilot experiment to demonstrate that Nested CRISPR can be carried out in a single injection through two sequential cuts by Cas9 and this finding would extend the advantages of Nested CRISPR to different animals and cellular systems.

## Materials and Methods

### Strains

We used the Bristol N2 strain as wild type background and worms were maintained on Nematode Growth Medium (NGM) plates seeded with *E. coli* OP50 bacteria (Stiernagle 2006). All strains generated in this study are listed in **Table S4**.

### crRNA and ssODN design

The 20-nucleotide guide sequences were selected with the help of CCTop (Stemmer *et al*. 2015) or Benchling (*www.benchling.com*), which are CRISPR/Cas9 target predictors. The crRNAs were ordered as 2 nmol products from IDT (*www.idtdna.com*) and were resuspended in 20 μl of nuclease-free duplex buffer to yield a stock concentration of 100 μM. Once the cut site had been determined, ssODN donors were designed in such a way that the EGFP 1-3 or mCherry 1-3 sequences were inserted in-frame immediately before the stop codon of the gene of interest (*pgl*-*1* and *prpf*-*4*), or within a few amino acids before the stop codon (*gtbp*-*1* and *ubh*-*4*), depending on the availability of a PAM sequence. The canonical design is as follows: a 35- to 45-bp left homology arm extending towards the cut site at or before the stop codon, followed by the EGFP 1-3 or mCherry 1-3 sequence, followed by a 35- to 45-bp right homology arm extending towards the 3’ UTR. The exact lengths of the homology arms depend on the distance of the insertion from the cut site and must account for the adjustment of nucleotides to ensure that the EGFP or mCherry 1-3 fragment is inserted in-frame. In the case of transcriptional reporters (K12C11.3), two crRNAs were designed to cut within the gene of interest, leaving behind a few amino acids after the start codon and before the stop codon. However, nucleotides encoding for these excess amino acids were not included in the ssODN repair template, thus generating mCherry 1-3 insertions that are immediately flanked by the start and stop codons. ssODNs were ordered as 4 nmol ultramers from IDT and were resuspended in 40 μl of nuclease-free duplex buffer to yield a stock concentration of 100 μM. The sequences of crRNAs and ssODN donors used in all EGFP 1-3 or mCherry 1-3 injections are shown in **Table S1 and Table S2**.

### Preparation of EGFP or mCherry PCR product repair template

The plasmids pJJR82 and pJJR83 were gifts from Mike Boxem (Addgene plasmids #75027 and #75028, respectively). These plasmids contain sequences for the EGFP and mCherry fluorophores respectively, which are codon-optimized for *C. elegans.* Primers were designed to allow the amplification of the complete sequence of EGFP or mCherry using the Phusion High-Fidelity DNA Polymerase (Thermo Fisher Scientific). Primers and PCR conditions for these universal steps are specified in the supplementary materials (Figure S2). 5 μl of PCR product were run on a 2% agarose gel to verify correct amplification of the fragments. The amplicon lengths for EGFP and mCherry are 865 bp and 855 bp, respectively. Eight to twelve 50-μl reactions were then purified with the MinElute PCR purification kit (QIAGEN). The yield achieved was usually around 800 – 1200 ng/μL.

### Preparation of mix

The individual components were mixed in the following order: the tracrRNA, *dpy*-*10* crRNA, target gene crRNA (or in the case of step 2, EGFP 1-3 or mCherry 1-3 crRNA), and Cas9 (IDT, cat. no. 1081058) were combined and incubated at 37°C for 10 minutes. Cas9 was added to the mix at distinct concentrations, from 0.25 to 1.64 ng/μl (**Table 1**). The *dpy*-*10* repair template and target gene ssODN (in the case of step 1) or EGFP/mCherry PCR product (in the case of step 2) were added, and the volume brought up to the required amount with nuclease-free H_2_O. In our experience, it is possible to prepare 5-μl injection mixes to avoid wasting excess reagents. The mixture was then centrifuged at 13,000 rpm for 2 minutes to settle particulate matter and was kept on ice prior to loading the capillary needles. Fresh injection mixes were normally prepared in this study. However, we observed that excess mix can be stored at −20°C and reused for future injections for a period of at least 4 months. The recommended concentrations for each component of the injection mix are specified in the supplementary materials (**Figure S3**).

### Microinjection

Approximately 1 μl of the injection mix was loaded on Eppendorf Femtotips Microinjection Capillary Tips (Eppendorf) using Eppendorf Microloader Pipette Tips (Eppendorf). Approximately 15 to 20 young adult hermaphrodites were immobilized in 2% agar pads with halocarbon oil and were injected with the corresponding transformation mix using the *XenoWorks* Microinjection System (Sutter Instrument) and the Nikon eclipse Ti-s inverted microscope with *Nomarski* optics. Injected worms were recovered in M9 buffer and were individually separated onto NGM plates. The plates were incubated at 20°C for 4 days or at 25°C for 3 days.

### Screening

F_1_ rollers and dumpys were individually transferred onto NGM plates and were left to lay F_2_ progeny. Single-worm or pooled (2-3 individuals) PCR was then performed on F_1_ worms. Primers were designed for each target gene and amplicon size shifts on 2% agarose gel were indicative of insertion events. If the PCR product was of the correct size, eight wild-type appearing F_2_ progeny were individually transferred onto NGM plates to isolate homozygous individuals. PCR products for homozygous animals were then purified using the QIAquick PCR purification kit (QIAGEN) and underwent Sanger Sequencing to verify the correctness of the insertion. In step 2 insertions, visual screening can be performed through fluorescence microscopy, in addition to genotyping by single-worm PCR. Green (EGFP) or red (mCherry) fluorescence were indicative of complete, in-frame insertion events. A list of external primers used for genotyping are shown in **Table S5**.

## Acknowledgments

We thank Mike Boxem for sharing the plasmids. We also thank Denis Dupuy, Montserrat Porta-de-la-Riva, and Peter Askjaer for critical reading of the manuscript and members of the Cerón lab for their comments on this study. Thanks to @shookstudio for their assistance in making figures 1, 2A and 4, and to www.mariaceron.com for making the supplementary videos. This work has been supported by a grant from the Instituto de Salud Carlos III (ISCIII) to JC (PI15-00895), cofunded by FEDER funds/ European Regional Development Fund (ERDF) — a way to Build Europe. JV has an INPhINIT PhD fellowship from La Caixa Foundation, and XS has a FPU PhD fellowship from MINECO.

## Tables

**Table S1.**
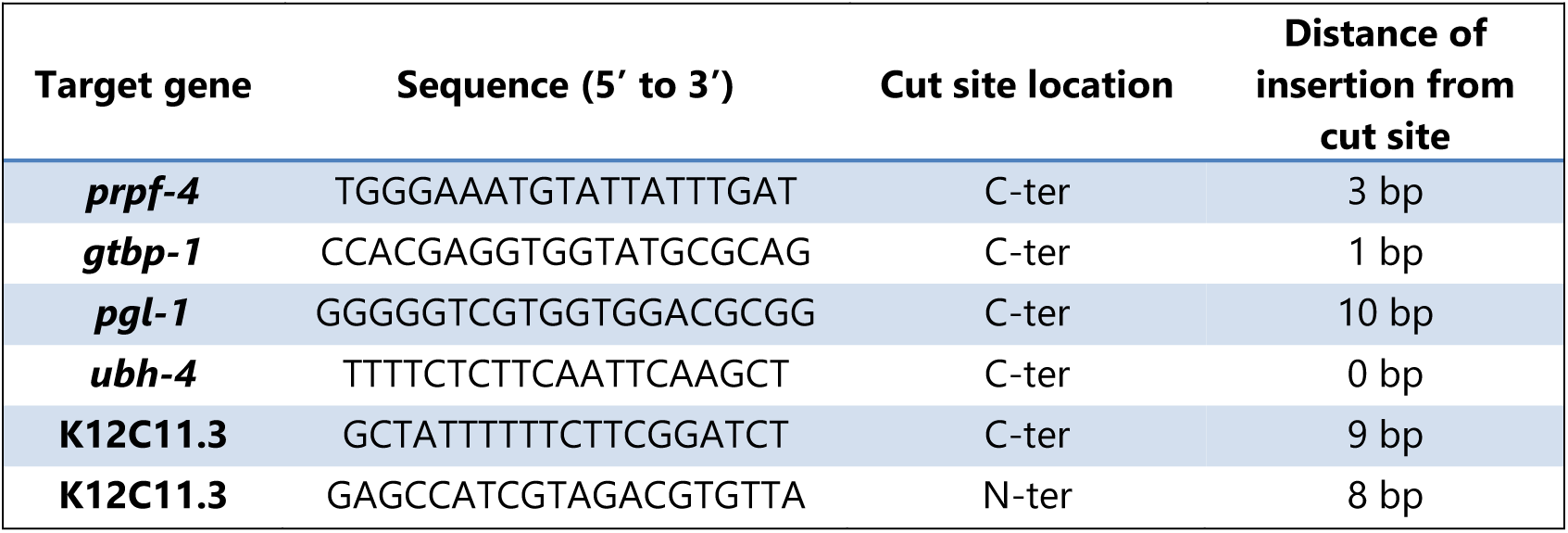
List of crRNAs used for Nested CRISPR Step 1.

**Table S2.**
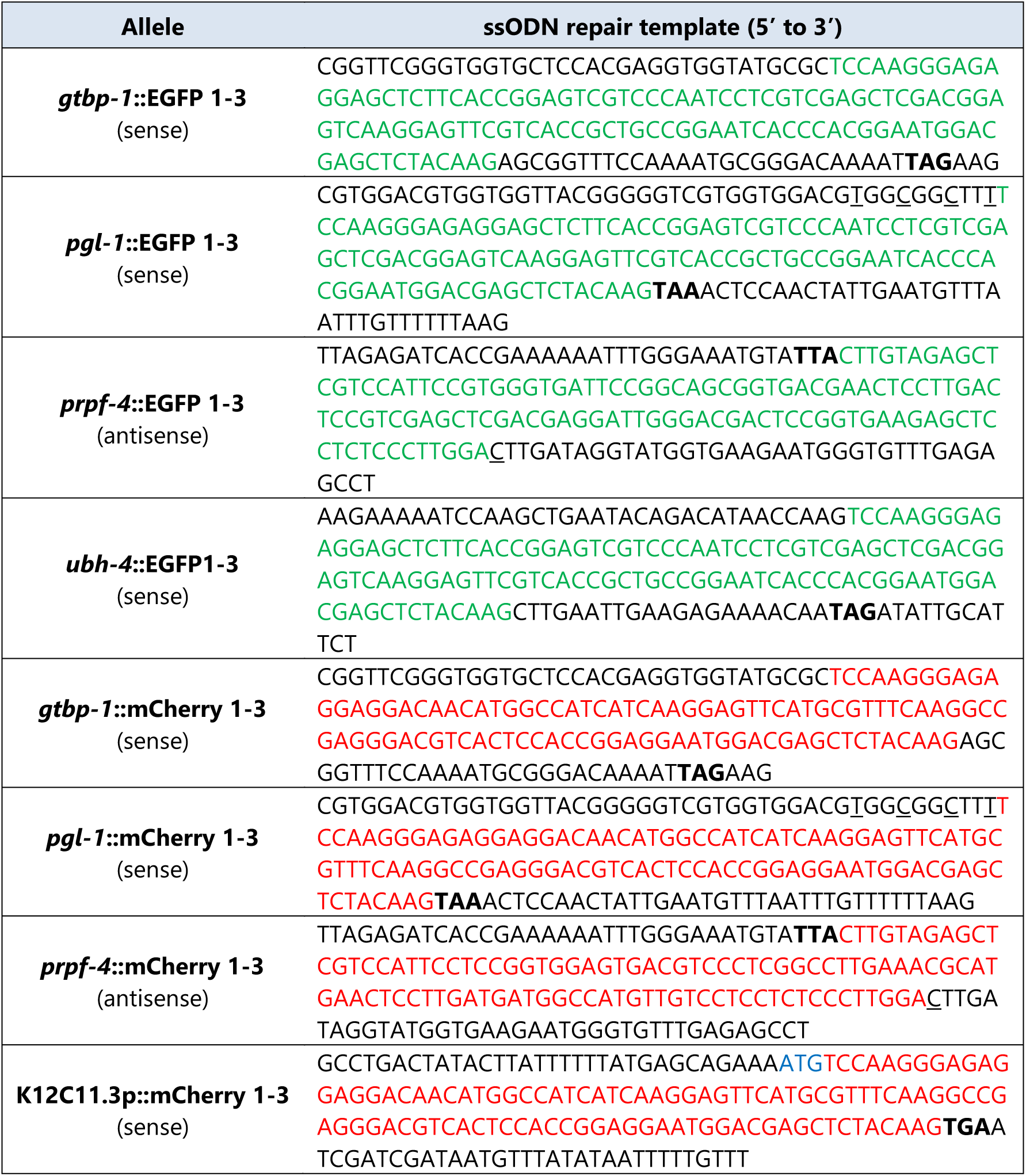
List of ssODNs used for Nested CRISPR Step 1. Red and green nucleotides represent EGFP 1-3 and mCherry 1-3 sequences, respectively. Nucleotides in blue and bold represent start and stop codons, respectively; and underscored nucleotides represent silent mutations.

**Table S3.**
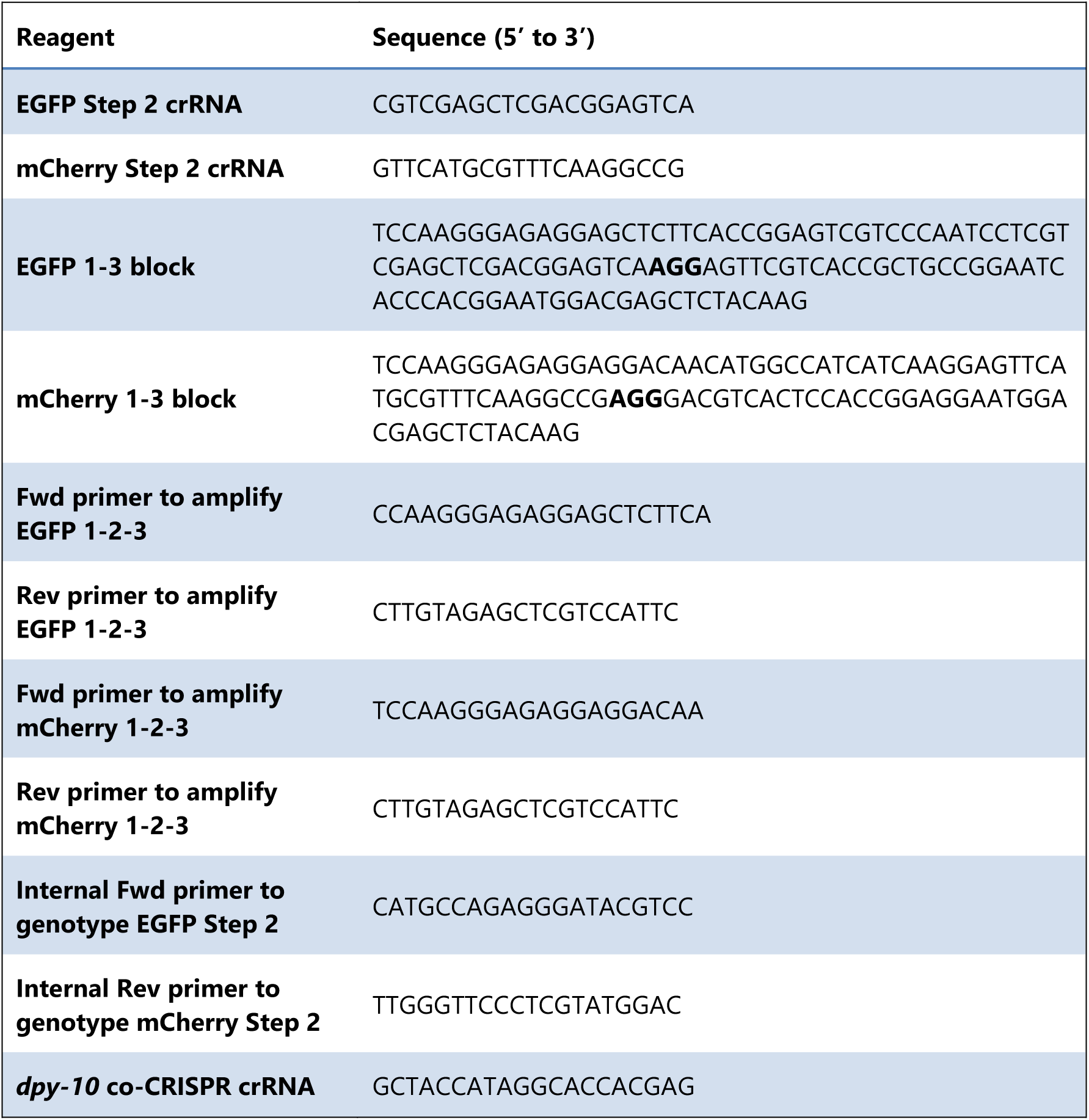
List of Nested CRISPR Universal Sequences. Nucleotides in bold represent PAM sequences.

**Table S4.**
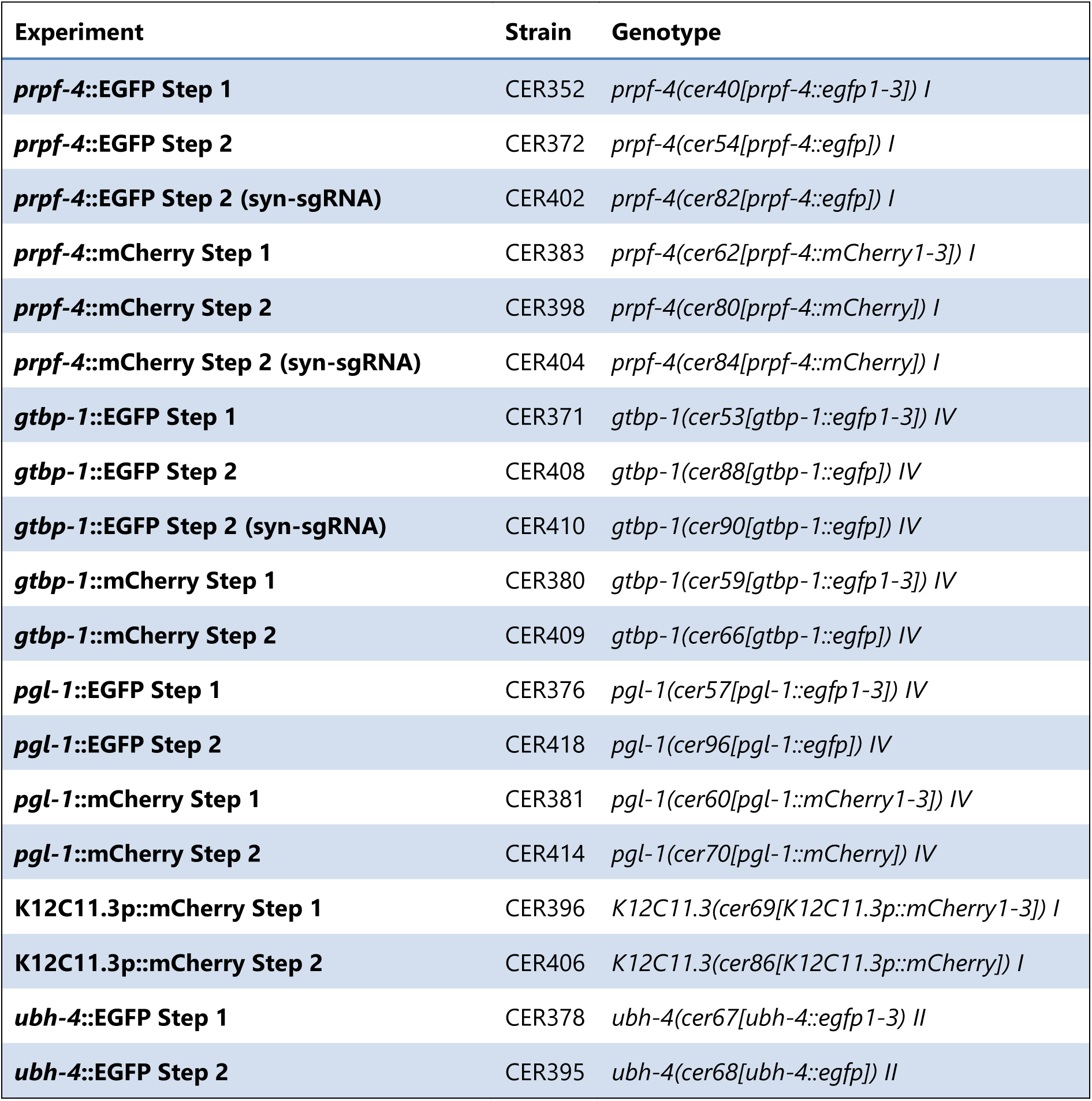
List of generated strains.

**Table S5.**
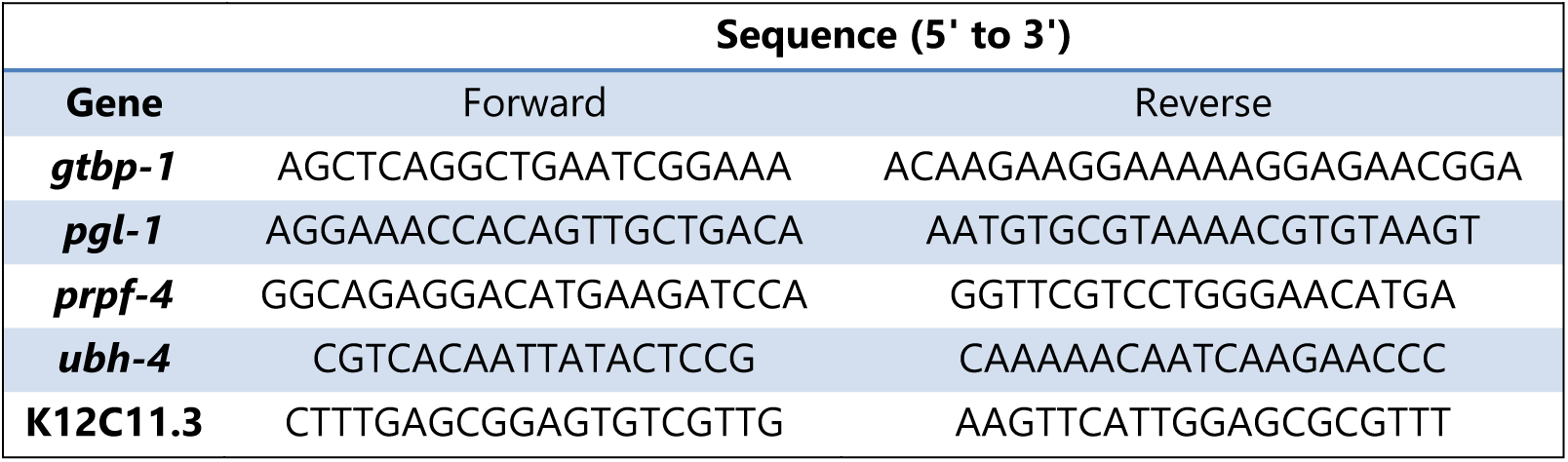
List of external primers for genotyping.

**Figure S1:**
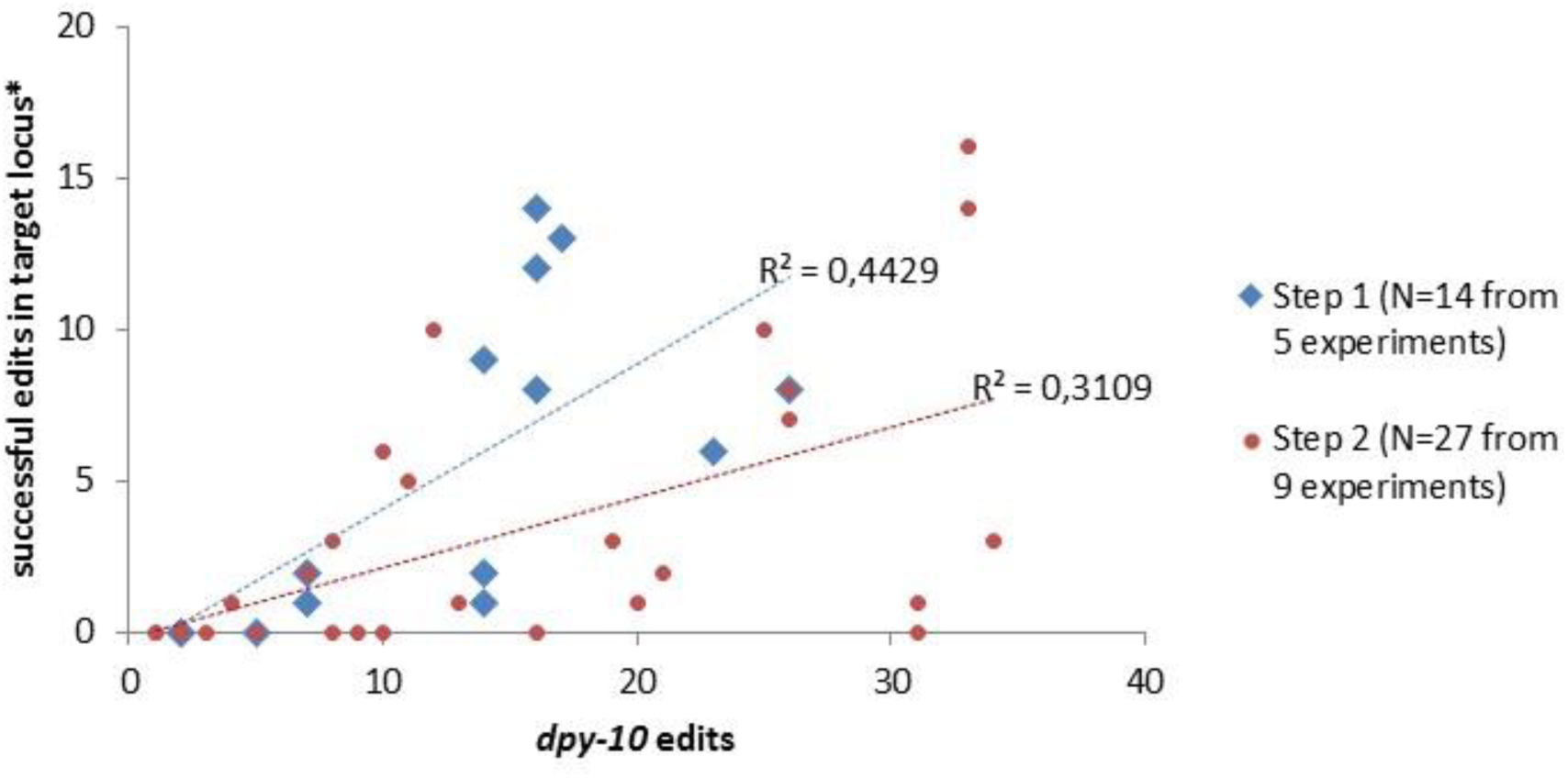
Correlation between the number of *dpy*-*10* edits and successful edits in the target locus. (^∗^based on bands of the correct size after screening by PCR). Each point represents an injected P0 worm giving rise to *dpy*-*10* edited F_1_ progeny. There is a moderate positive correlation between the number of *dpy*-*10* edited progeny and the number of successful edits in both step 1 and step 2 experiments, with step 1 experiments having a slightly higher correlation. This demonstrates that a high number of *dpy*-*10* edits does not necessarily correspond to higher editing efficiencies in the target locus.

**Figure S2.**
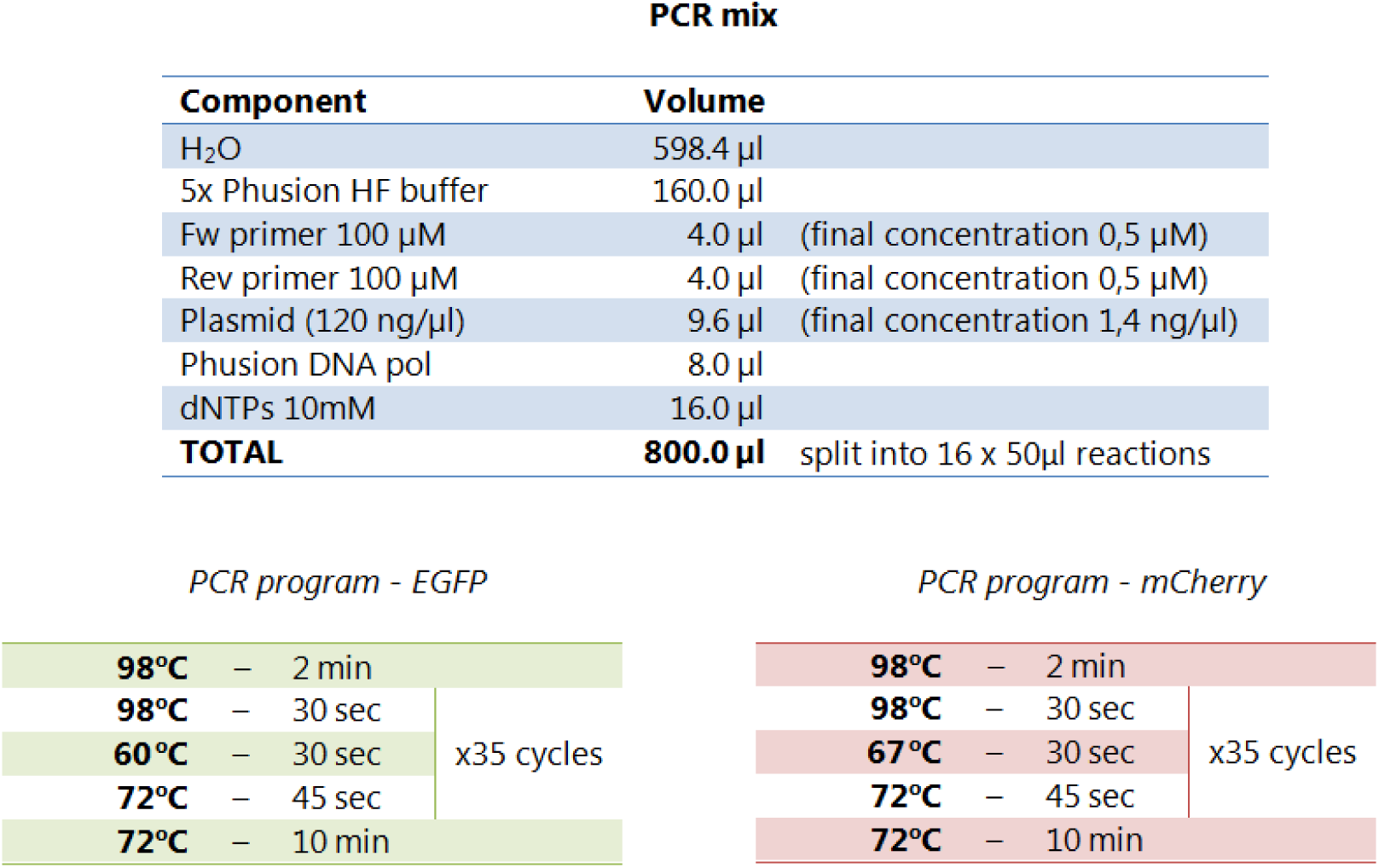
Reagents and conditions for generation of universal PCR product repair template for Nested CRISPR Step 2.

**Figure S3.**
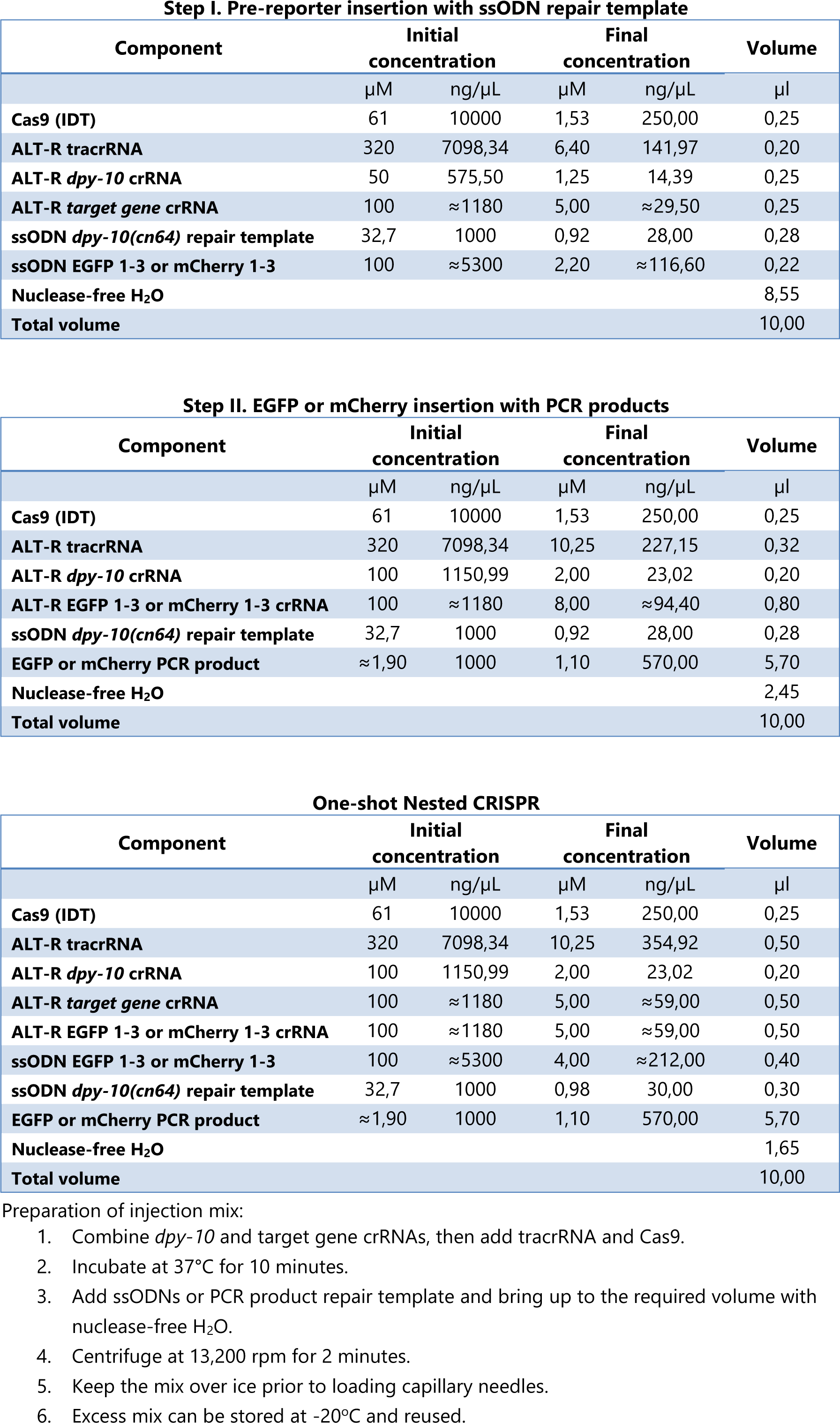
Standard composition of injection mixes for Nested CRISPR.

## References

Arribere J. A., R. T. Bell, B. X. H. Fu, K. L. Artiles, P. S. Hartman, et al., 2014 Efficient marker-free recovery of custom genetic modifications with CRISPR/Cas9 in Caenorhabditis elegans. Genetics 198: 837–846. https://doi.org/10.1534/genetics.114.169730

Bell R. T., B. X. H. Fu, and A. Z. Fire, 2016 Cas9 Variants Expand the Target Repertoire in Caenorhabditis elegans. Genetics 202: 381–388. https://doi.org/10.1534/genetics.115.185041

Briner A. E., P. D. Donohoue, A. A. Gomaa, K. Selle, E. M. Slorach, et al., 2014 Guide RNA functional modules direct Cas9 activity and orthogonality. Mol. Cell 56: 333–339. https://doi.org/10.1016/j.molcel.2014.09.019

Chen C., L. A. Fenk, and M. de Bono, 2013 Efficient genome editing in Caenorhabditis elegans by CRISPR-targeted homologous recombination. Nucleic Acids Res. 41: e193. https://doi.org/10.1093/nar/gkt805

Chen X., F. Xu, C. Zhu, J. Ji, X. Zhou, et al., 2014 Dual sgRNA-directed gene knockout using CRISPR/Cas9 technology in Caenorhabditis elegans. Sci. Rep. 4: 7581. https://doi.org/10.1038/srep07581

Cho S. W., J. Lee, D. Carroll, J.-S. Kim, and J. Lee, 2013 Heritable gene knockout in Caenorhabditis elegans by direct injection of Cas9-sgRNA ribonucleoproteins. Genetics 195: 1177–1180. https://doi.org/10.1534/genetics.113.155853

Dickinson D. J., J. D. Ward, D. J. Reiner, and B. Goldstein, 2013 Engineering the Caenorhabditis elegans genome using Cas9-triggered homologous recombination. Nat. Methods 10: 1028–1034. https://doi.org/10.1038/nmeth.2641

Dokshin G. A., K. S. Ghanta, K. M. Piscopo, and C. C. Mello, 2018 Robust Genome Editing With Short Single-Stranded and Long, Partially Single-Stranded DNA Donors in Caenorhabditiselegans. Genetics. https://doi.org/10.1534/genetics.118.301532

Farboud B., and B. J. Meyer, 2015 Dramatic enhancement of genome editing by CRISPR/Cas9 through improved guide RNA design. Genetics 199: 959–971. https://doi.org/10.1534/genetics.115.175166

Frokjaer-Jensen C., 2013 Exciting prospects for precise engineering of Caenorhabditis elegans genomes with CRISPR/Cas9. Genetics 195: 635–642. https://doi.org/10.1534/genetics.113.156521

Ghanta K. S., G. A. Dokshin, A. Mir, P. M. Krishnamurthy, H. Gneid, et al., 2018 5’ Modifications Improve Potency and Efficacy of DNA Donors for Precision Genome Editing. bioRxiv.

Gurumurthy C., R. Quadros, J. Adams, P. Alcaide, S. Ayabe, et al., 2018 Re-Evaluating One-step Generation of Mice Carrying Conditional Alleles by CRISPR-Cas9-Mediated Genome Editing Technology. bioRxiv.

Heyn P., A. T. Kalinka, P. Tomancak, and K. M. Neugebauer, 2015 Introns and gene expression: cellular constraints, transcriptional regulation, and evolutionary consequences. Bioessays 37: 148–154. https://doi.org/10.1002/bies.201400138

Jinek M., K. Chylinski, I. Fonfara, M. Hauer, and J. A. Doudna, 2012 A Programmable Dual-RNA – Guided DNA Endonuclease in Adaptive Bacterial Immunity. Science (80-.). 337: 816–821.

Katic I., L. Xu, and R. Ciosk, 2015 CRISPR/Cas9 Genome Editing in Caenorhabditis elegans: Evaluation of Templates for Homology-Mediated Repair and Knock-Ins by Homology-Independent DNA Repair. G3 (Bethesda). 5: 1649–1656. https://doi.org/10.1534/g3.115.019273

Kemphues K., 2005 Essential genes. WormBook 1–7. https://doi.org/10.1895/wormbook.1.57.1

Kim H., T. Ishidate, K. S. Ghanta, M. Seth, D. J. Conte, et al., 2014 A co-CRISPR strategy for efficient genome editing in Caenorhabditis elegans. Genetics 197: 1069–1080. https://doi.org/10.1534/genetics.114.166389

Lim Y., S. Y. Bak, K. Sung, E. Jeong, S. H. Lee, et al., 2016 Structural roles of guide RNAs in the nuclease activity of Cas9 endonuclease. Nat. Commun. 7: 13350. https://doi.org/10.1038/ncomms13350

Liu W., S. Li, Y. Zhang, J. Li, and Y. Li, 2018 Efficient CRISPR-based genome editing using tandem guide RNAs and editable surrogate reporters. FEBS Open Bio 8: 1167–1175. https://doi.org/10.1002/2211-5463.12437

McDiarmid T. A., V. Au, A. Loewen, J. J. H. Liang, K. Mizumoto, et al., 2018 CRISPR-Cas9 human gene replacement and phenomic characterization in *Caenorhabditis elegans* to understand the functional conservation of human genes and decipher variants of uncertain significance. bioRxiv.

Mojica F. J. M., C. Diez-Villasenor, J. Garcia-Martinez, and E. Soria, 2005 Intervening sequences of regularly spaced prokaryotic repeats derive from foreign genetic elements. J. Mol. Evol. 60: 174–182. https://doi.org/10.1007/s00239-004-0046-3

Norris A. D., H.-M. Kim, M. P. Colaiacovo, and J. A. Calarco, 2015 Efficient Genome Editing in Caenorhabditis elegans with a Toolkit of Dual-Marker Selection Cassettes. Genetics 201: 449–458. https://doi.org/10.1534/genetics.115.180679

Paix A., Y. Wang, H. E. Smith, C.-Y. S. Lee, D. Calidas, et al., 2014 Scalable and versatile genome editing using linear DNAs with microhomology to Cas9 Sites in Caenorhabditis elegans. Genetics 198: 1347–1356. https://doi.org/10.1534/genetics.114.170423

Paix A., A. Folkmann, D. Rasoloson, and G. Seydoux, 2015 High Efficiency, Homology-Directed Genome Editing in Caenorhabditis elegans Using CRISPR-Cas9 Ribonucleoprotein Complexes. Genetics 201: 47–54. https://doi.org/10.1534/genetics.115.179382

Paix A., H. Schmidt, and G. Seydoux, 2016 Cas9-assisted recombineering in C. elegans: genome editing using in vivo assembly of linear DNAs. Nucleic Acids Res. 44: e128. https://doi.org/10.1093/nar/gkw502

Prior H., A. K. Jawad, L. MacConnachie, and A. A. Beg, 2017 Highly Efficient, Rapid and Co-CRISPR-Independent Genome Editing in Caenorhabditis elegans. G3 (Bethesda). 7: 3693–3698. https://doi.org/10.1534/g3.117.300216

Quadros R. M., H. Miura, D. W. Harms, H. Akatsuka, T. Sato, et al., 2017 Easi-CRISPR: a robust method for one-step generation of mice carrying conditional and insertion alleles using long ssDNA donors and CRISPR ribonucleoproteins. Genome Biol. 18: 92. https://doi.org/10.1186/s13059-017-1220-4

Ran F. A., P. D. Hsu, J. Wright, V. Agarwala, D. A. Scott, et al., 2013 Genome engineering using the CRISPR-Cas9 system. Nat. Protoc. 8: 2281–2308. https://doi.org/10.1038/nprot.2013.143

Schwartz M. L., and E. M. Jorgensen, 2016 SapTrap, a Toolkit for High-Throughput CRISPR/Cas9 Gene Modification in Caenorhabditis elegans. Genetics 202: 1277–1288. https://doi.org/10.1534/genetics.115.184275

Stemmer M., T. Thumberger, M. Del Sol Keyer, J. Wittbrodt, and J. L. Mateo, 2015 CCTop: An Intuitive, Flexible and Reliable CRISPR/Cas9 Target Prediction Tool. PLoS One 10: e0124633. https://doi.org/10.1371/journal.pone.0124633

Stiernagle T., 2006 Maintenance of C. elegans. WormBook 1–11. https://doi.org/10.1895/wormbook.1.101.1

Verkuijl S. A., and M. G. Rots, 2018 The influence of eukaryotic chromatin state on CRISPR-Cas9 editing efficiencies. Curr. Opin. Biotechnol. 55: 68–73. https://doi.org/10.1016/j.copbio.2018.07.005

Waaijers S., V. Portegijs, J. Kerver, B. B. L. G. Lemmens, M. Tijsterman, et al., 2013 CRISPR/Cas9-targeted mutagenesis in Caenorhabditis elegans. Genetics 195: 1187–1191. https://doi.org/10.1534/genetics.113.156299

Wang H., H. Park, J. Liu, and P. W. Sternberg, 2018 An efficient genome editing strategy to generate putative null mutants in Caenorhabditis elegans using CRISPR/Cas9. G3 (Bethesda). 8.

Yang H., H. Wang, C. S. Shivalila, A. W. Cheng, L. Shi, et al., 2013 One-step generation of mice carrying reporter and conditional alleles by CRISPR/Cas-mediated genome engineering. Cell 154: 1370–1379. https://doi.org/10.1016/j.cell.2013.08.022

